# Mapping and quantifying nascent transcript start sites using TT-TSS-seq

**DOI:** 10.1101/2025.03.25.645230

**Authors:** Eleanor Elgood Hunt, Claudia Vivori, Folkert J van Werven

## Abstract

Transcription initiation is a highly dynamic and tightly regulated process involving the coordinated action of transcription factors, chromatin remodelers, and RNA polymerase which determine where and when transcription begins. Accurately mapping and quantifying transcription start sites (TSSs) from nascently transcribed RNAs remains a key area of interest, as it provides critical insights into transcription dynamics. Here, we combined transient transcriptome sequencing with transcription start site sequencing (TT-TSS-seq) to accurately map and quantify transcription initiation sites from nascent transcripts. Since transient metabolic labelling yields low-input RNA, we optimized the TSS-seq protocol to enhance sensitivity and accuracy. Specifically, we refined enzymatic reactions for decapping and RNA ligation and incorporated 5’ oligonucleotides containing unique molecular identifiers (UMIs) and barcodes to enable accurate quantification and sample multiplexing. The TT-TSS-seq approach detected transcription initiation of unstable transcripts, such as enhancer RNAs. Moreover, we identified that a large fraction of genes use multiple transcription initiation sites, yet often produce only a single stable transcript. Overall, TT-TSS-seq provides precise mapping and quantification of transcription initiation sites, offering new insights into transcriptional dynamics and expanding the toolkit for studying gene regulation.

## Introduction

Transcription initiation is a critical regulatory step in which RNA polymerase II (RNAPII), guided by transcription factors and co-factors, is recruited to the promoter region and starts synthesizing RNA. The selection of transcription start sites (TSSs) plays a fundamental role in determining transcript diversity and gene expression patterns. Most gene loci harbour multiple TSSs, which are differentially regulated in response to cellular conditions, developmental cues, and environmental signals (Wang et al. 2008; Carninci et al. 2006; Lu and Lin 2019; Minghao Chia et al. 2021; Brown et al. 2014). Despite the prevalence of alternative TSS usage, the functional significance of this regulatory mechanism remains largely unknown. Understanding how transcription initiation is controlled and how different TSSs contribute to gene expression and cellular function is essential for uncovering new layers of gene regulation. Moreover, dysregulation of TSS selection has been linked to various diseases, including cancer and neurodevelopmental disorders, highlighting the need for further investigation into its underlying mechanisms (Thorsen et al. 2011; Demircioglu et al. 2019).

For mapping TSSs, sequencing approaches have been employed with different biochemical strategies to capture the 5’ ends of transcripts. Cap-trapping techniques, including CAGE (Cap Analysis of Gene Expression), are the most commonly used. This technique involves chemical treatment to oxidize the 5’ caps of RNA, enabling biotinylation and subsequent streptavidin pulldown to enrich for capped transcripts (Takahashi et al. 2012). Template-switching reverse transcription (TSRT), which involves reverse transcription using an enzyme which adds 1-3 non-templated bases (typically cytosine) at the 5’ end, works well for low-input material reactions (Policastro et al. 2020). Oligo-capping methods provide a high resolution but require a relatively large amount of input material. These processes involve the dephosphorylation of uncapped RNA, followed by enzymatic decapping to generate 5’ monophosphate ends, which are essential for subsequent adaptor ligation (Pelechano, Wei, and Steinmetz 2013; Arribere and Gilbert 2013).

Most TSS-based sequencing techniques use mature RNA, which enables the detection of TSSs of stably expressed transcripts but not of TSSs from unstable RNAs, such as long noncoding transcripts. Furthermore, using mature RNA prevents insights into transcription initiation dynamics. Therefore, nascent or newly transcribed RNA is required for detecting TSSs of unstable transcripts and quantitative analysis of transcription initiation. A few approaches have been developed to map and quantify TSS usage for nascent or newly synthesized RNA. For example in NET-CAGE, RNA still associated with chromatin is selectively captured, enabling TSS mapping of actively transcribed genes (Hirabayashi et al. 2019). Isolation of RNAPII-associated nascent RNAs (e.g. POINT-5-seq), instead, provides insights into transcription initiation and early elongation (Sousa-Luis et al. 2021). *In vitro* run-on assays such as GRO-cap and PRO-cap label and sequence newly synthesized RNA to determine TSSs (Danko et al. 2015; Mahat et al. 2016). Short-capped RNA sequencing (scaRNA-seq) selectively captures short, capped transcripts to identify sites of transcription initiation (Larke et al. 2021). Limitations of some of these approaches include the need for complex biochemical purifications or fractionations and the risk of contamination with steady-state RNAs.

Here, we combined transient metabolic labelling with the oligo-capping approach to map sites of transcription initiation at the nucleotide resolution (TT-TSS-seq). We optimized the chemical and enzymatic reaction conditions for TSS-seq and applied it to transiently-labelled RNA isolated from mouse embryonic stem cells (mESCs) (Gregersen, Mitter, and Svejstrup 2020; Minghao Chia et al. 2021). We show that the TT-TSS-seq approach enables precise mapping of TSSs from nascent RNAs including unstable transcripts. TT-TSS-seq is highly reproducible, through *in vivo* labeling of unperturbed cells (without requiring nuclear isolation or *in vitro* reactions) is relatively simple and, serves as a powerful tool for studying the dynamics of transcription initiation.

## Results and Discussion

### Optimization of TSS-seq protocol

To develop TSS-seq for nascent or newly transcribed RNA, we improved the previously described TSS-seq protocol for relatively low RNA input (Figure 1A) (Minghao Chia et al. 2021). We chose an oligo-capping strategy because it enables the detection of TSSs at single-nucleotide resolution (Pelechano, Wei, and Steinmetz 2013; Minghao Chia et al. 2021). In order to optimize the protocol for low amounts of RNA, we incorporated part of the iCLIP2 protocol for library preparation (Buchbender et al. 2020). First, cells are pulsed with 4-thiouridine (4SU) resulting in the labelling of newly synthesised RNA. RNA is treated with alkaline phosphatase to remove the 5’ phosphate groups from non-capped RNA. Next, the mRNA decapping enzyme (MDE) removes the 5’-terminal caps, exposing the 5’ phosphate group exclusively on once-capped transcripts. Subsequently, a 5’ RNA-DNA hybrid adaptor containing a barcode for sample multiplexing and a unique molecular identifier (UMI) is ligated. After the pooling of samples, polyadenylated (polyA+) RNA or transiently 4SU labelled RNA populations are purified (for polyA-TSS-seq or TT-TSS-seq, respectively). RNA is fragmented, the 3’ end fixed, before a pre-adenylated 3’ adaptor is ligated using a truncated T4 RNA ligase that can only utilise pre-adenylated substrates. RNA is then reverse-transcribed and the cDNA pre-amplified before the first size selection is carried out to remove primer dimers and short inserts. A second amplification and size selection are performed, followed by quality control and sequencing.

**Figure 1.**
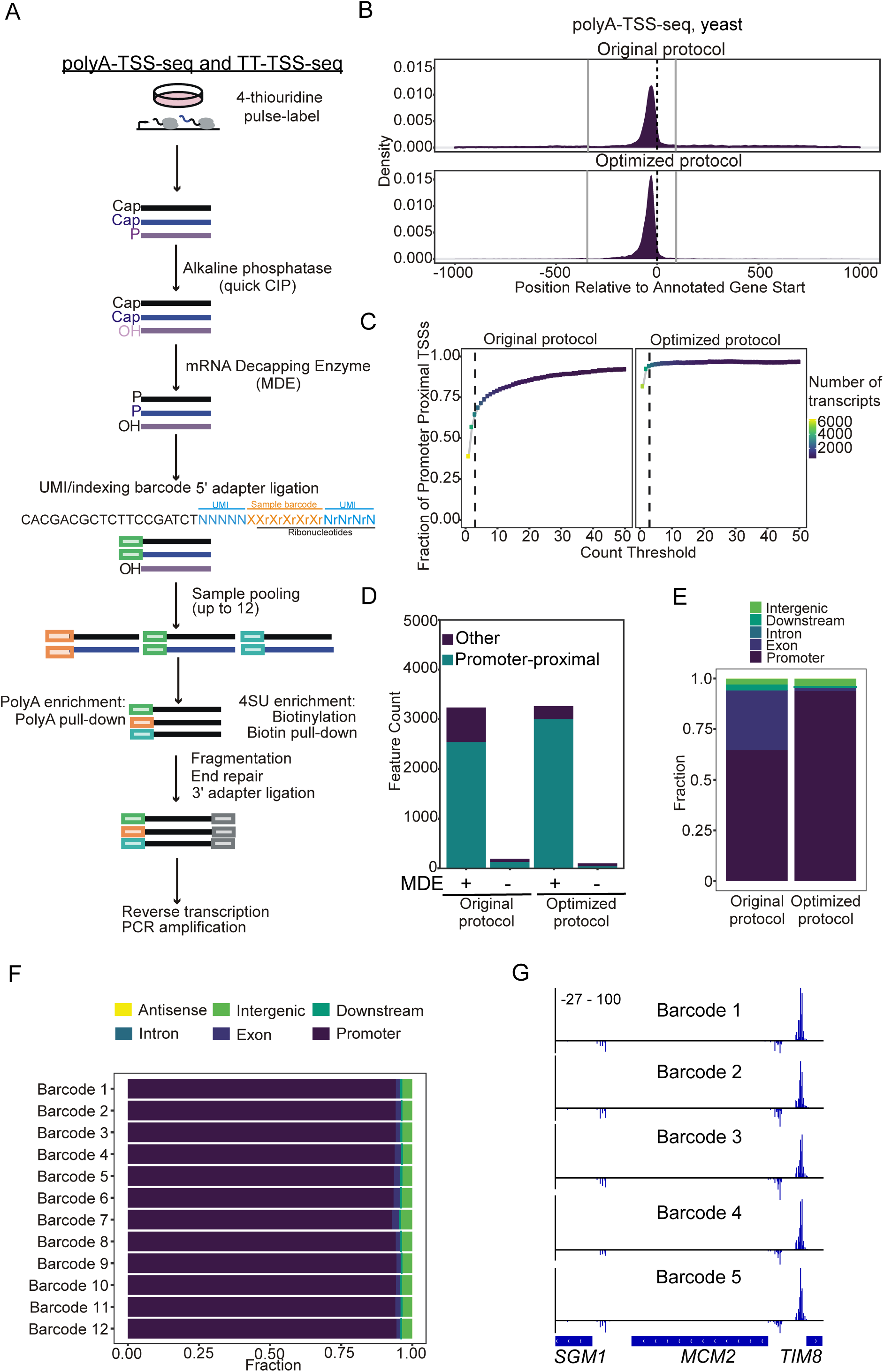
Optimization of TSS-seq protocol with yeast RNA. **(A)** Schematic of the TSS-sequencing protocol. Alkaline phosphatase (quickCIP) removes the 5’ phosphate groups from non-capped fragments (shown in purple). The mRNA decapping enzyme (MDE) removes the 5’-terminal caps exposing a 5’phosphate group (shown in black). This enables the ligation of a 5’ adaptor, which contains sample barcodes to allow sample multiplexing and UMIs. Nascent or steady-state RNA is then selected, before fragmentation, end-repair, 3’ adaptor ligation, reverse transcription and PCR amplification. **(B)** Density plot showing tag locations compared to annotated gene start codons for polyA-TSS-seq. RNA isolated from yeast was subjected to polyA-TSS-seq using either the original or optimised conditions. The x-axis is centred on the annotated open reading frame (ORF) start codon. n=2 biological repeats. **(C)** Fraction (y-axis) of tags located in promoter-proximal regions (−300 to +100 bp from annotated ORF start codons) at different read thresholds (x-axis). **(D)** Bar plot showing the number of tags located in promoter-proximal regions with the original or optimized polyA-TSS-seq protocols (+ MDE). As negative controls, samples that had not undergone the decapping reaction were included (−MDE). A threshold of n=3 counts was applied. **(E)** Genomic locations of detected tags. Tag counts were normalised using the DESeq2 median-of-ratios approach using a threshold of n=3 counts. (F) Genomic locations of detected TSSs using the different 5’ adaptor barcode sequences (Document S1). **(G)** Integrated genome browser (IGV) showing an example locus of TSS-seq signals for barcodes 1 to 6.

Multiple steps of the TSS-seq protocol were optimised, including fragmentation, reverse transcription, RNA clean-up, PCR and DNA size selection conditions. In addition, we increased the efficiency of the alkaline phosphatase, MDE, and ligations reactions (Table S1). We denatured RNA before ligation to decrease any effects of RNA structure altering the reaction time, temperature and polyethylene glycol (PEG) concentration.

To compare the optimized protocol to the original one, we first performed polyA-TSS-seq on RNA purified from yeast cells. The 5’-most nucleotides, referred to as “tags”, of the reads were extracted for mapping of TSSs. The optimized protocol showed an increased fraction of the tags located in the promoter-proximal region (−300 to +100 bp from annotated open reading frame start) across different count thresholds (Figure 1B and 1C). Additionally, the optimized protocol showed an increase in the number of tags (threshold of 3 or more) in the promoter-proximal region whilst decreasing the number of detected tags in the negative control, compared to the tags detected in the original conditions (Figure 1D). TSS-seq with the optimized protocol showed an increase in the fraction of tags in the promoter-proximal region (94% vs. 65% with the original protocol) (Figure 1E). In conclusion, the optimized TSS-seq protocol increased the accuracy of TSS detection in polyA+ RNA isolated from yeast. We recently applied the optimized TSS-seq promoter to study how DNA supercoiling affects TSS usage (reference to be included).

### Multiplexing of samples

In the optimized TSS-seq protocol, we incorporated a barcode into the 5’ adaptor to enable sample multiplexing. To assess whether the barcoded 5’ adaptors gave comparable tag profiles, we carried out polyA-TSS-seq with yeast RNA. Dephosphorylation and decapping were carried out in a single reaction before samples were split and 12 different barcoded 5’ adaptors were ligated. These samples were then pooled together and PCR was performed using primers with the same index.

The polyA-TSS-seq samples generated with barcoded 5’ adaptors showed a comparable genomic distribution (Figure 1F). Libraries produced using barcoded 5’ adaptors showed a good correlation with each other (Pearson’s r=0.92-0.98) (Figure S1A). This suggests that the different barcodes did not introduce more significant variations or biases in ligation. We determined the dinucleotide frequencies to assess whether there was a bias in adapter ligation. The first nucleotide represents the −1 nucleotide and the second nucleotide represents the +1 tag/TSS identified by polyA-TSS-seq (Figure S1B). We observed enrichment for +1 As in the tags/TSSs and no clear differences in dinucleotide usage between the barcodes, which is consistent with previous findings (Policastro et al. 2020). This suggested that the presence of the barcode does not bias towards the ligation of a specific 5’ nucleotide. Thus, multiplexing samples, as described for the optimized TSS-seq protocol, is a feasible strategy for handling multiple samples and may offer potential advantages in reducing sample loss in the subsequent steps of the protocol.

To further validate the optimised polyA-TSS-seq protocol, it was similarly applied to RNA isolated from mESCs. The identified tags showed strong enrichment to annotated TSSs (Figure 2A). For further analysis, we defined the promoter-proximal region as −500 to +500 bp from annotated TSSs. The proportion of promoter-proximal tags was determined at different count thresholds and more than 90% of tags (using a threshold of 3 counts) were annotated to promoter-proximal regions (Figure 2B and 2C). Furthermore, approximately 31,500 transcripts featured at minimum one TSS (Figure 2D). We conclude that the optimized polyA-TSS-seq protocol is suitable for more complex transcriptomes.

**Figure 2.**
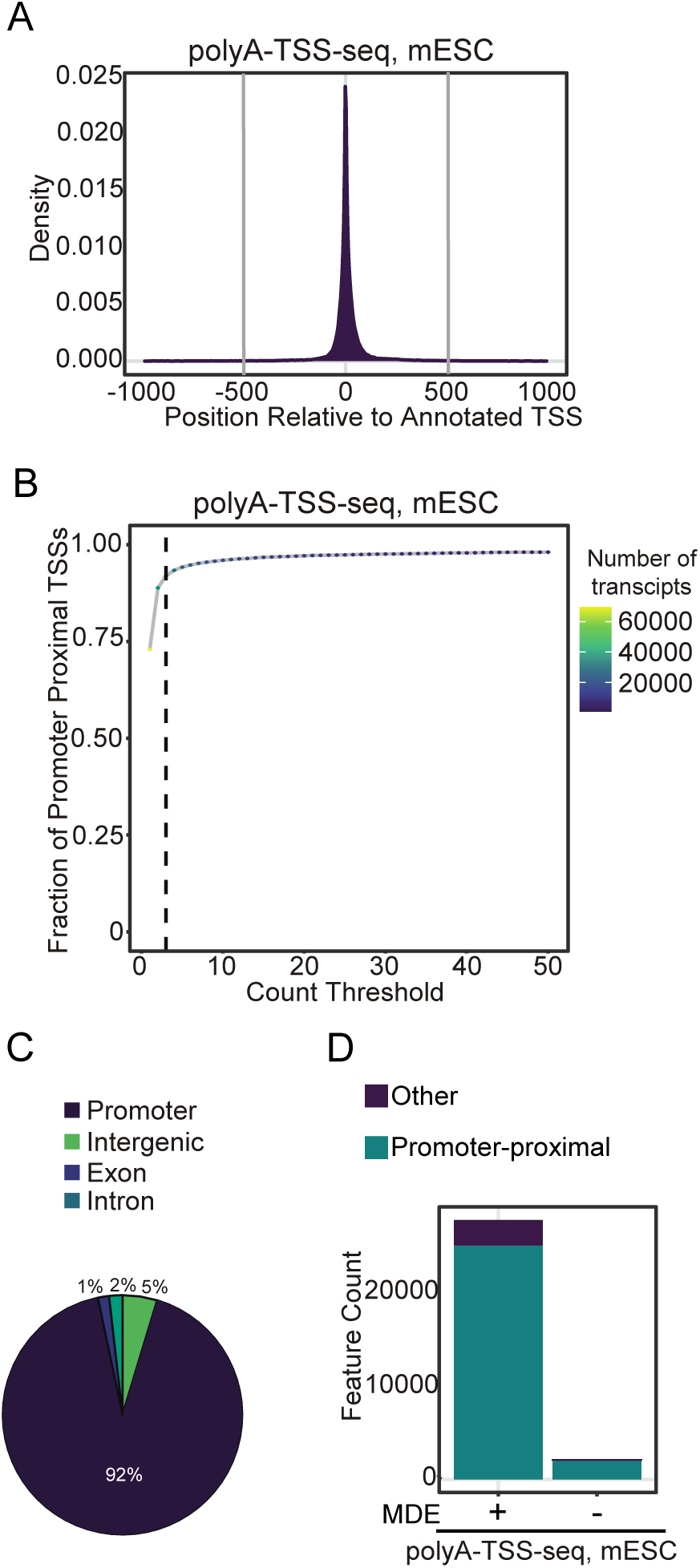
Optimized TSS-seq protocol tested with mESC RNA. **(A)** Density plot showing tag locations detected with polyA-TSS-seq using mESC RNA compared to annotated mouse TSSs. n=2 biological repeats. **(B)** Fraction of tags that are promoter-proximal (−500 to +500 bp from annotated TSSs) at different read count thresholds for mESC. A threshold of n=3 read counts was used. **(C)** Genomic locations of detected tags with polyA-TSS-seq. Tag counts were normalised using the DESeq2 median-of-ratios approach using a threshold n=3 counts. **(D)** Bar plot showing the number of transcripts with promoter-proximal tags in mESCs detected by polyA-TSS-seq (+MDE). As a negative control, a no-decapping reaction sample was included (−MDE). A threshold of n=3 counts was applied.

### Transient 4SU labelling of RNA combined with TSS-seq (TT-TSS-seq)

Multiple methods have been described to enrich nascent or newly transcribed RNAs (Hirabayashi et al. 2019; Kruesi et al. 2013; Larke et al. 2021; Sousa-Luis et al. 2021). Transient labelling of RNA with 4SU provides a way to isolate nascent transcripts, with advantages including high reproducibility and *in vivo* labelling of unperturbed cells (TT-seq or TT-chem-seq) (Gregersen, Mitter, and Svejstrup 2020; Schwalb et al. 2016). TT-seq methods have been widely used to study transcription dynamics. To balance RNA yield without compromising transcription dynamics, we assessed the degree of 4SU incorporation following different labelling times and using different 4SU concentrations in mESCs (Figure S2A). With increased labelling time or 4SU concentration, increased 4SU incorporation occurred, as expected, and no signal was detected in the control (no 4SU) (Figure S2A). Similar to the TT-chem-seq protocol, we used 15 minutes of labelling with 1mM 4SU for further analysis (Gregersen, Mitter, and Svejstrup 2020).

We carried out TSS-seq on purified RNA that was transiently labelled with 4SU (TT-TSS-seq). For comparison, we also used another method to enrich nascent RNA and selected transcripts with less than 300 nucleotides, as in scaRNA-seq (Larke et al. 2021) (Figure S2B). We compared the TT-TSS-seq profile to the profiles of polyA-TSS-seq and scaRNA-seq (Figure 3A) (Larke et al. 2021; Henriques et al. 2018; Nechaev et al. 2010). The polyA-TSS-seq and TT-TSS-seq detected tags showed strong enrichment at the annotated TSSs (Figure 3B). In contrast, the tags detected with scaRNA-seq also showed signals downstream of annotated TSSs suggesting the detection of RNA decay intermediates (Figure 3B). For each sample, the regions with multiple TSSs were clustered into transcript start regions (TSRs), and the correlation between TSR signals detected by each method was assessed. The TSR signals showed a good correlation between biological replicates (Pearson’s r=0.86 for polyA-TSS-seq, 0.89 for TT-TSS-seq, and 0.96 for scaRNA-seq) (Figure S2C). However, TSRs detected using scaRNA-seq showed little-to-no correlation with TT-TSS-seq (Pearson’s r=0-0.22), while TT-TSS-seq and polyA-TSS-seq showed some correlation (Pearson’s r=0.41-0.58).

**Figure 3.**
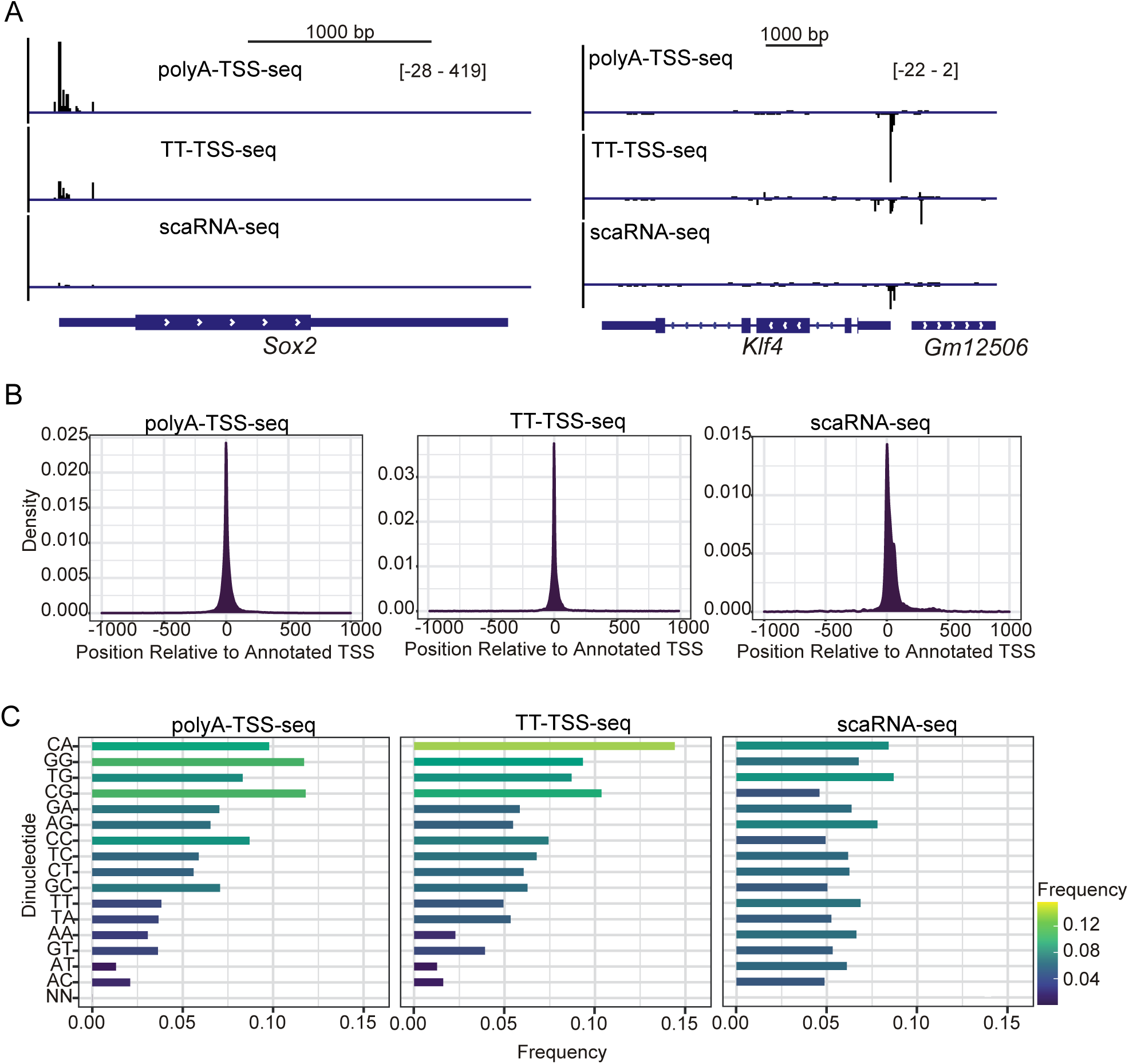
TT-TSS-seq protocol applied on RNA isolated from mESCs. **(A)** Integrated genome viewer (IGV) showing example loci (Sox2 and Klf4) of TSSs detected using mESC polyA-TSS-seq, TT-TSS-seq and scaRNA-seq (less than 300 nt). Scale is indicated on the top right of each panel. (B) Density plots showing TSS locations detected using mESC polyA-TSS-seq, TT-TSS-seq and scaRNA-seq (less than 300 nt) compared to annotated mouse TSSs. n=2 biological repeats. **(C)** Dinucleotide frequencies analysis. The first nucleotide represents the −1 nucleotide and the second nucleotide represents the +1 tag/TSS identified by polyA-TSS-seq. A threshold of n=3 counts was used.

We also assessed the dinucleotide frequency in the detected TSRs. A preference for pyrimidines at the −1 and purines at the +1 nucleotide (the TSS) has previously been identified (Vo Ngoc et al. 2017). The tags detected by polyA-TSS-seq on RNA isolated from mESCs showed enrichment for guanine (Figure 3C). This is also consistent with the more GC-rich nature of mammalian promoters (Policastro et al. 2020). TSSs identified with TT-TSS-seq showed an increased preference for CA_+1_ and a decreased preference for GG_+1_ and CG_+1_ compared to polyA-TSS-seq (Figure 3C). This is in line with other nascent TSS-sequencing techniques (such as GRO-Cap), that identified a preference for +1 A (Vo Ngoc et al. 2017; Luse et al. 2020). Similarly, GRO-Cap identified a decreased preference for +1 G and C TSSs in unstable transcripts compared to stable ones (Core et al. 2014). By contrast, the TSSs detected here using scaRNA-seq did not show a preference for a dinucleotide combination (Figure 3C).

We conclude that TT-TSS-seq is suitable for mapping and quantifying transcription initiation sites. The TSSs detected by scaRNA-seq showed poor correlation with TSSs detected using TT-TSS-seq and poly-TSS-seq and showed a lack of dinucleotide preference. A possible explanation is that other RNA species are present in the scaRNA-seq samples. For example, RNA degradation intermediates may be enriched during size selection, which would create more substrates for the alkaline phosphatase reaction, potentially resulting in incomplete dephosphorylation.

### TT-TSS-seq detects TSSs of unstable RNAs

The TSSs identified using the TT-TSS-seq protocol showed less enrichment in the promoter-proximal region, however, it is unclear whether this reflects technical limitations or limitations in current TSS annotations (Figure 4A). Therefore, we further investigated the differences between the polyA-TSS-seq and TT-TSS-seq datasets. As before, the TSSs were clustered into TSRs, and the number of transcripts with detected TSRs was determined. The number of identified transcripts with TSRs was increased by using TT-TSS-seq compared to polyA-TSS-seq (Figure 4B). Next, we carried out differential analysis on the TSRs detected by TT-TSS-seq and polyA-TSS-seq (Figure 4D). Approximately 11,500 TSRs were detected at significantly increased levels and approximately 2,500 at significantly decreased levels in TT-TSS-seq compared to polyA-TSS-seq. More transcripts with retained introns were associated with TSRs detected at significantly higher levels in TT-TSS-seq compared to polyA-TSS-seq (Figure 4D).

**Figure 4.**
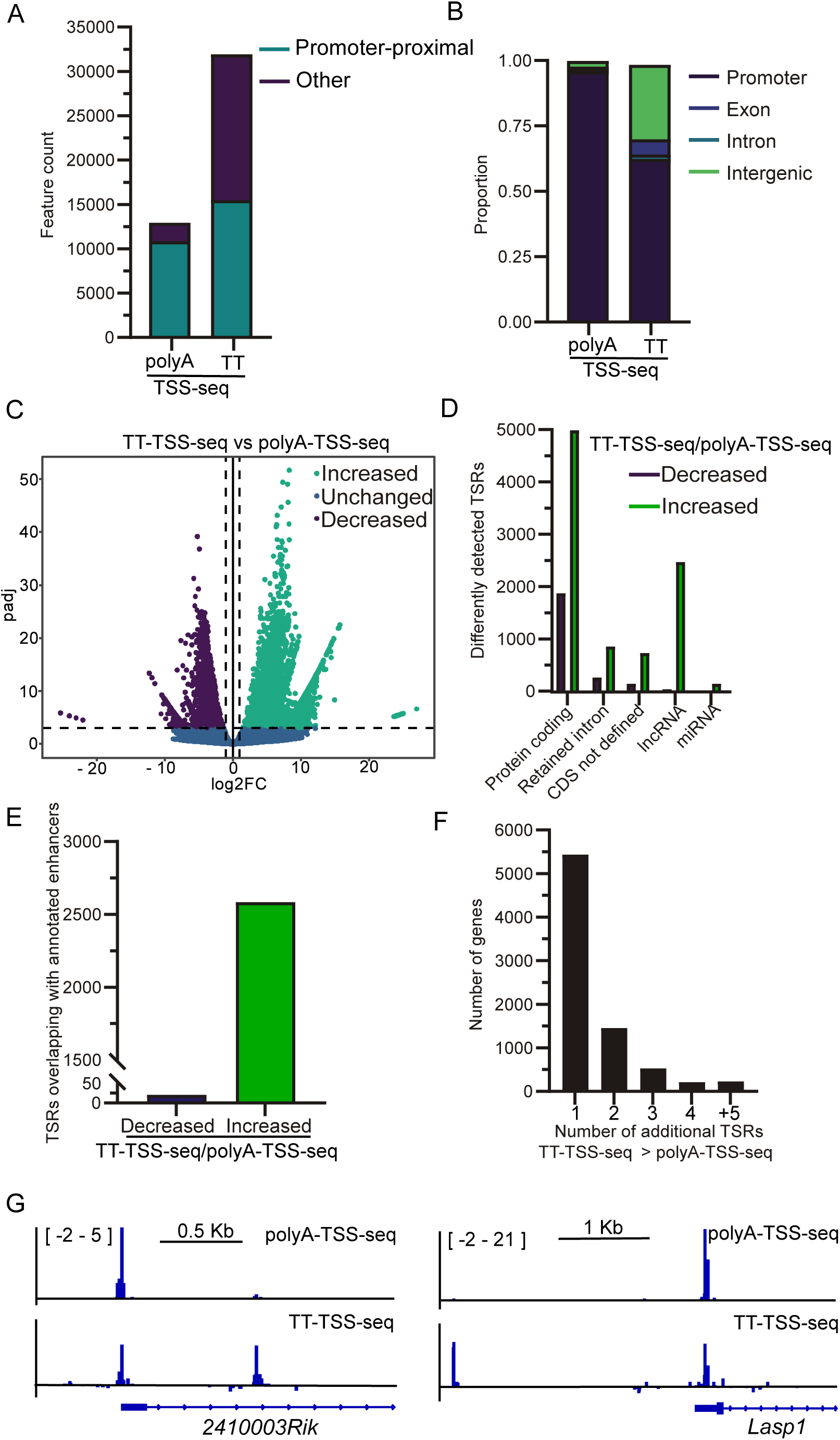
Comparing tags identified with polyA-TSS-seq and TT-TSS-seq. **(A)** Bar plot showing the number of transcripts with promoter-proximal TSRs. Normalised tags within 25 bp, and to a maximum distance of 250 bp, were merged to identify TSRs. **(B)** Bar graph showing the distribution of detected TSSs amongst genic regions. −500 to +500 bp from annotated TSSs was used to define the promoter-proximal region. UMI-based PCR duplicate removal was carried out during TSS-sequencing analysis. Tag counts were normalised using the DESeq2 median-of-ratios approach using a threshold of 3 counts. **(C)** Volcano plot showing the TSRs detected at significantly different levels when using nascent compared to steady-state RNA. Normalised TSSs, located within 25 bp and to a maximum distance of 250 bp were grouped into TSRs and differential analysis was carried out (using DESeq2 with a false discovery threshold of 0.001 and a fold-change threshold of 2). **(D)** Number of TSRs, differentially detected between TT-TSS-seq versus polyA-TSS-seq associated with different transcript types (as annotated by Ensembl). “Protein-coding transcripts” harbour ORFs, “CDS not defined” are alternatively spliced isoforms of protein-coding transcripts with no identified ORF, and “retained intron transcripts” are alternatively spliced isoforms that harbour an intron and are predicted to be non-coding. **(E)** Number of TSRs, differentially detected between TT-TSS-seq and polyA-TSS-seq that overlap with enhancer regions (as annotated by FANTOM5) (Noguchi et al. 2017). **(F)** The number of genes with additional TSRs detected with a significantly increased signal detected by TT-TSS-seq compared to polyA-TSS-seq. Genes with a single TSR detected in the polyA-TSS-seq, which did not show a significant difference in signal or exhibited an increase compared to TT-TSS-seq, were used for the analysis. The number of additional TSRs detected at significantly higher levels by TT-TSS-seq is displayed. **(G)** IGV showing example loci of TT-TSS-seq and polyA-TSS-seq. Scale is indicated on the top.

Next, we assessed whether the differentially detected TSRs overlapped with enhancer regions. Over 2,500 TSRs detected at significantly higher levels using TT-TSS-seq overlapped with annotated enhancer regions (Figure 4E). Indeed, it is known that most enhancer RNAs contain a 5’cap (Sartorelli and Lauberth 2020). Finally, we determined whether using TT-TSS-seq increased the detection of alternative TSSs. We selected genes with one TSR that showed no significant difference in signal or were increased in polyA-TSS-seq compared to TT-TSS-seq. We found that nearly 8,000 genes had at least one additional TSR detected at significantly higher levels by TT-TSS-seq (Figure 4F and 4G).

In summary, the TT-TSS-seq protocol effectively identifies a broad spectrum of TSSs, including alternative TSSs. Notably, TT-TSS-seq detects short-lived RNA species such as enhancer RNAs, long-noncoding RNAs, and transcripts predicted to undergo rapid decay. These findings highlight the sensitivity of TT-TSS-seq in capturing transcription initiation events with high accuracy. We propose that TT-TSS-seq is a powerful and reliable method for mapping transcription initiation sites across the mammalian genome, providing valuable insights into gene regulation and transcriptome dynamics.

## Methods

### Yeast growth conditions

*S. cerevisiae* BY4741 strain background (derived from S288C) was used for the analysis of the yeast part of the manuscript. Cells were grown in liquid cultures (YPD (1.0% (w/v) yeast extract, 2.0% (w/v) peptone, 2.0% (w/v) glucose, and supplemented with uracil (2.4 mg/l) and adenine (1.2 mg/l))) in incubator/shaker (30°C, 300 rpm) till the exponential phase.

### mESCs culturing

mESCs (HM1) were grown in 2i/LIF [49% DMEM/F12-/- (Gibco), 49% Neurobasal (Gibco), 1X N2 (Gibco), 1X B27 (Gibco), 0.05 mM 2-Mercaptoethanol (Gibco), 2mM L-Glutamine (Gibco), Recombinant mouse 50,000 units LIF (ESGRO), 1 µM PD03259010 (Stemgent), 3 µM CHIR 99021 (TOCRIS), 1% Penicillin/Streptomycin]]. Cell culture surfaces were coated with 0.15% gelatin in PBS, for a minimum of 10 minutes at 37°C for mESC culture. mESCs were maintained at 37°C, 5% CO2, and confirmed to be negative for mycoplasma. mESCs were passed every 2-3 days depending on colony size, with media change occurring in the intermediate days. To pass cells, one wash with PBS was performed, before incubating the cells with Accutase (Gibco) at 37°C for exactly 3 minutes.

### RNA extraction

RNA was extracted from yeast as previously detailed (M. Chia et al. 2021). For RNA extraction from yeast grown in YPD, 24 optical density (OD) units for TSS-seq were collected by centrifugation, and snap-frozen in liquid nitrogen. Per 10 OD units, RNA was extracted with 1 ml Tris-EDTA-SDS (TES) buffer (10 mM Tris-HCl pH 7.5, 10 mM EDTA, 0.5% SDS) and 1 ml Acid Phenol:Chloroform:Isoamyl alcohol (125:24:1, Ambion), by shaking at 65°C for 45 minutes at 1400 rpm. After centrifugation at 4°C, max speed, 10 minutes, the aqueous phase was transferred to cold ethanol with 0.3 M sodium acetate. Precipitation was carried out overnight at 4°C, before centrifugation at 4°C, max speed, 20 minutes. The pellet was washed once with 80% v/v ethanol, dried, and reconstituted in DEPC-treated sterile water. rDNase (Machery-Nagel) treatment was carried out for 20 minutes at 37°C, before spin column purification (Machery-Nagel).

RNA was extracted from mammalian cells using TRIzol (Thermo Fisher). For TRIzol RNA extraction, 1 ml of TRIzol was added per 10-cm dish. Chloroform (0.2 volume, Thermo Fisher) was added, and the mixture was shaken for 30 seconds before centrifugation was performed. The aqueous phase was mixed with 1.1 volumes of isopropanol (Thermo Fisher), and RNA was precipitated overnight at 4°C. After centrifugation, the pellet was washed with 85% (v/v) ethanol, dried, and reconstituted in DEPC-treated sterile water. rDNase (Qiagen) digestion was carried out at 37°C for 1 hour before purification by spin column (Qiagen) or by phenol:chloroform and ethanol precipitation.

For small RNA size selection, Monarch RNA Cleanup with a 1:3 ratio of sample: ethanol volume was used.

### 4SU RNA labelling and assessment

4SU labelling was carried out as detailed previously (Gregersen, Mitter, and Svejstrup 2020), with few modifications. Cell culture media was removed, filtered, and 1 mM 4SU (Glentham Life Sciences) was added. The cells were returned to the incubator, and labelled with 4SU for 15 minutes before medium aspiration and TRIzol addition. 1/5 volume of chloroform was added, mixed, and then centrifuged at 12,000g for 15 minutes at 4°C. 1.1 volumes of isopropanol and 1 µl of glycoblue were mixed with the upper aqueous layer, precipitation then occurred overnight at −20°C. The pellet was then washed once with 85% ethanol, dried, and resuspended.

RNA integrity was checked for a selection of samples using Bioanalyser (Agilent Technologies). 4SU incorporation was assessed using slot blot. Here, 5 µg of total RNA was mixed with 3 µl biotin buffer (833 mM Tris-HCl, pH 7.4, and 83.3 mM EDTA) and 50 µl of 0.1 mg/ml MTSEA biotin-XX linker (Biotium, dissolved in dimethylformamide; Sigma-Aldrich) to a total volume of 200 µl, and incubated in the dark for 30 minutes. RNA clean-up was performed by adding an equal volume of phenol/chloroform/isoamyl alcohol (25:24:1 (vol/vol/vol)), mixing, and centrifuging at 12,000g for 5 minutes at 4°C. The aqueous phase was mixed with 1.1 volumes of isopropanol, 1/10 volume of 5 M NaCl and 1 µl of glycoblue and precipitated at −20°C for at least 2 hours. The pellet was washed once with 85% ethanol, dried, and resuspended to 10 µl.

Samples were dropped onto the Hybond-N-membrane in the slot blot apparatus before UV crosslinking was performed using 0.2 J/cm2 (254 nm) in a Stratalinker. The membrane was blocked in blocking buffer (10% (w/v) SDS, 1 mM EDTA, PBS) for 20 minutes, before being incubated with HRP-conjugated streptavidin (1:50,000 dilution of 1 mg/ml) in blocking buffer for 15 minutes at room temperature. The membrane was then washed 2 times in blocking buffer, 2 times in wash buffer 1 (1% (w/v) SDS, PBS), and 2 times in wash buffer 2 (0.1% (w/v) SDS, PBS), for 10 minutes for each wash. Streptavidin-HRP signal was visualised using Enhanced Chemiluminescence (ECL) reagent. RNA levels were assessed by staining with 0.5 M sodium acetate and 0.5% w/v methylene blue for 10 minutes, before multiple washes with water.

### TSS-seq library preparation

The optimized TSS-seq protocol was adapted from previously described protocols (Arribere and Gilbert 2013; Malabat et al. 2015; Pelechano, Wei, and Steinmetz 2013; Minghao Chia et al. 2021). The library preparation steps were adapted, as described, for the iCLIP2 protocol (Buchbender et al. 2020). DNase digestion was performed on total RNA (DNase I, Qiagen, for 1 hour at 37°C). Following DNase treatment, RNA was purified using phenol/chloroform/isoamyl alcohol and precipitated overnight at −20°C using 1.1 volumes of isopropanol, 1/10 volume of 5 M NaCl and 1 µl of glycoblue. At least 69 µg total RNA was taken per sample.

Total RNA was dephosphorylated with quickCIP (NEB, 1.2 U/µg RNA) for 2 hours, 37°C, with RNasin Plus before heat inactivation of the enzyme at 80°C for 2 minutes. RNA was extracted using phenol/chloroform/isoamyl alcohol as before. RNA was treated with mRNA decapping enzyme (NEB, 0.7 U/µg RNA) for 2 hours, 37°C with RNasin Plus. For the negative control, no decapping enzyme was included. RNA was purified using phenol/chloroform/isoamyl alcohol as before. RNA and the 5’ adaptor (10 µM) were denatured for 2 minutes at 70°C, then cooled on ice for 2 minutes. Ligation was performed for 2 hours, 25°C then 16 hours, 16°C using T4 RNA ligase I (30 U) with RNasin Plus. RNAClean XP (1.8X ratio, Beckman Coulter) was used to remove unligated adaptors, according to the manufacturer’s instructions. The concentration of the samples was measured using Qubit RNA BR assay, and samples were multiplexed. Samples were split according to the expected yield (3% of polyA+ RNA and 0.7% of 4SU+ RNA). polyA+ RNA was enriched using Dynabeads Oligo(dT)25 (Thermo Fisher) or 4SU+ RNA was purified as detailed in the TT-chem-seq protocol (Gregersen, Mitter, and Svejstrup 2020). Briefly, RNA was biotinylated as above, and reconstituted to 50 µl. 200 µl of μMACS streptavidin MicroBeads (Miltenyi) were added and mixed to the biotinylated RNA on a rotating wheel for 20 minutes at room temperature. μColumns were equilibrated with nucleic acid equilibration buffer before the samples were applied, and the flow-through non-4SU labelled RNA was collected. Columns were washed twice with 55°C wash buffer (100 mM Tris-HCl, pH 7.4, 10 mM EDTA, 1 M NaCl and 0.1% v/v Tween 20) before 4SU positive RNA was eluted by addition twice of 100 mM DTT (Sigma-Aldrich). Flow-through and eluted fractions were cleaned up using RNeasy MinElute kits (Qiagen).

Yeast RNA was fragmented for 3 min 15 sec and mammalian RNA for 5 min 15 sec at 70°C with alkaline fragmentation reagent (Ambion), to achieve 200-300 bp fragments. In scaRNA-seq samples, RNA fragmentation was not performed. RNeasy MinElute Clean-up columns, with 1.5x volume of Ethanol, were used to purify the samples. The 3’ ends were fixed by CIP treatment (NEB, 30 U), with RNasin, at 37°C for 1 hour. Heat inactivation was carried out at 80°C for 2 min before the RNA was extracted using phenol/chloroform/isoamyl alcohol as before. RNA, PEG8000 (10%) and pre-adenylated 3’ adaptor (2 µM) were denatured for 2 minutes at 70°C, then cooled on ice for 2 minutes. Ligation was carried out by T4 Rnl2tr (NEB,200 U) for 2 hours, 25°C then 16 hours, 16°C. RNAClean XP (1.8X ratio) was used to remove unligated adaptors. RNA, RT oligo (0.5 pmol) and dNTP (10 mM each) were denatured for 5 minutes at 65°C, then cooled on ice for 1 minute. Reverse transcription was carried out by SuperScript III (2 µl), with RNasin Plus with the following conditions: 25°C for 5 minutes, 42°C for 20 minutes, 50°C for 40 minutes, 80°C for 5 minutes. Template RNA was removed by RNase H (10 U) for 37°C, 30 minutes. AMPure XP beads (Agencourt, Beckman Coulter) were used to clean up the cDNA. 3x beads and 1.7x isopropanol added to the reverse transcription reaction and incubated for 5 minutes. Beads were washed twice with 85% ethanol, dried, and eluted twice in nuclease-free water. cDNA pre-amplification was performed with i5_s and i7_s (300 nM each) primers and Phusion HF PCR Mastermix (Thermo Fisher). 6 PCR cycles were carried out (98°C for 10 seconds, 65°C for 30 seconds, 72°C for 30 seconds, with 3 minutes at 72°C final extension). ProNex beads (Promega) with a 1:2.95 ratio to sample were used to remove sequences less than 55 nt (including primer dimers). Final amplification was carried out with NEBNext i50 and i70 primers (500 nM each) and Phusion HF PCR Mastermix, for 8 cycles. Amplification was checked by using Novex 6% TBE (Tris/Borate/EDTA) gel and staining with SYBR green. ProNex beads (Promega) with a 1:2.3 ratio to sample were used for purification. Library concentration was measured using Qubit High Sensitivity D1000. Libraries were sequenced with Illumina NovaSeq 6000 using 100 nt paired-end reads, with 50 million reads minimum per sample, at the Genomics Facility of the Francis Crick Institute.

### Mapping of TSS-seq reads

Read demultiplexing was performed using Ultraplex with parameters “--phredquality 15 −-min_length 0” (Wilkins et al. 2021). Adaptor trimming was performed using Cutadapt with parameters “--minimum-length 20” (Martin 2011). Bowtie2 was used for pre-mapping to ribosomal and small RNAs (Langmead and Salzberg 2012). Genome mapping was performed using STAR, for mouse samples with Ensembl GRCm38 (mm10) release-89 annotation, and for S288C samples with S. cerevisiae Ensembl R64-1-1 release-90 annotation (Dobin et al. 2013). Umitools was used for deduplication (Smith, Heger, and Sudbery 2017). Bedtools was used to identify the 5’-most nucleotide (tag) and for genome-wide comparative analyses (Quinlan and Hall 2010).

### Data analysis TSS-seq

The majority of downstream analysis was performed using TSRexploreR and CAGEr (Policastro et al. 2021; Haberle et al. 2015). The promoter-proximal region was defined as −300 to +100 bp from annotated yeast start codons (based on *S. cerevisiae* Ensembl R64-1-1 release-90 annotation for S288C), or −500 to +500 bp from annotated mouse TSSs (based on GRCm38 (mm10) release-89 annotation).

For TSRexploreR analysis, a 5’ tag threshold of 3 was used. DESeq2 median-of-ratios approach was used for normalisation. For mammalian TSS analysis, the 5’-most tags were grouped into transcript start regions (TSRs) using a maximum distance of 25 bp and maximum total width of 250 bp. Pearson’s correlation between samples was calculated using the normalised counts of each TSR. DESeq2 was used to identify differentially expressed TSRs, using log2(fold-change)>=1, and false discovery rate (FDR)<0.001 (Love, Huber, and Anders 2014). Transcript biotype annotation from Ensembl release 112 was used (Harrison et al. 2024). Overlap between identified TSRs and CAGE-annotated enhancer regions was carried out using the GRanges package (Noguchi et al. 2017). TSRs were associated with the closest gene on the same strand, giving priority to the promoter annotation, and the number of genes with multiple associated TSRs was calculated.

### Statistical analysis

Information regarding any statistical tests used, number of samples, or number of biological replicate experiments is stated in the corresponding figure legends.

## Publicly available datasets used in this study

CAGE data were obtained from (Noguchi et al. 2017).

## Data Availability

The accession number for the TSS sequencing data reported in this paper is GSE292786.

## Supplementary documents

Table S1.

Figure S1.

Figure S2.

Document S1.

## Acknowledgements

We thank the Crick Genomics STP for sequencing the libraries. We also thank the members of the van Werven lab for the critical reading of the manuscript. This work was supported by the Francis Crick Institute (CC2043), which receives its core funding from Cancer Research UK (CC2043), the UK Medical Research Council (CC2043), and the Wellcome Trust (CC2043).

## References

Arribere, Joshua A., and Wendy V. Gilbert. 2013. “Roles for Transcript Leaders in Translation and MRNA Decay Revealed by Transcript Leader Sequencing.” Genome Research 23 (6): 977–87.

Brown, J. B., N. Boley, R. Eisman, G. E. May, M. H. Stoiber, M. O. Duff, B. W. Booth, et al. 2014. “Diversity and Dynamics of the Drosophila Transcriptome.” Nature 512 (7515): 393–99.

Buchbender, A., H. Mutter, F. X. R. Sutandy, N. Kortel, H. Hanel, A. Busch, S. Ebersberger, and J. Konig. 2020. “Improved Library Preparation with the New ICLIP2 Protocol.” Methods 178: 33–48.

Carninci, Piero, Albin Sandelin, Boris Lenhard, Shintaro Katayama, Kazuro Shimokawa, Jasmina Ponjavic, Colin A. M. Semple, et al. 2006. “Genome-Wide Analysis of Mammalian Promoter Architecture and Evolution.” Nature Genetics 38 (6): 626–35.

Chia, M., C. Li, S. Marques, V. Pelechano, N. M. Luscombe, and F. J. van Werven. 2021. “High-Resolution Analysis of Cell-State Transitions in Yeast Suggests Widespread Transcriptional Tuning by Alternative Starts.” Genome Biology 22 (1): 34.

Chia, Minghao, Cai Li, Sueli Marques, Vicente Pelechano, Nicholas M. Luscombe, and Folkert J. van Werven. 2021. “Author Correction: High-Resolution Analysis of Cell-State Transitions in Yeast Suggests Widespread Transcriptional Tuning by Alternative Starts.” Genome Biology 22 (1): 46.

Core, Leighton J., André L. Martins, Charles G. Danko, Colin T. Waters, Adam Siepel, and John T. Lis. 2014. “Analysis of Nascent RNA Identifies a Unified Architecture of Initiation Regions at Mammalian Promoters and Enhancers.” Nature Genetics 46 (12): 1311–20.

Danko, Charles G., Stephanie L. Hyland, Leighton J. Core, Andre L. Martins, Colin T. Waters, Hyung Won Lee, Vivian G. Cheung, W. Lee Kraus, John T. Lis, and Adam Siepel. 2015. “Identification of Active Transcriptional Regulatory Elements from GRO-Seq Data.” Nature Methods 12 (5): 433–38.

Demircioglu, D., E. Cukuroglu, M. Kindermans, T. Nandi, C. Calabrese, N. A. Fonseca, A. Kahles, et al. 2019. “A Pan-Cancer Transcriptome Analysis Reveals Pervasive Regulation through Alternative Promoters.” Cell 178 (6): 1465–1477 e17.

Dobin, Alexander, Carrie A. Davis, Felix Schlesinger, Jorg Drenkow, Chris Zaleski, Sonali Jha, Philippe Batut, Mark Chaisson, and Thomas R. Gingeras. 2013. “STAR: Ultrafast Universal RNA-Seq Aligner.” Bioinformatics (Oxford, England) 29 (1): 15–21.

Gregersen, Lea H., Richard Mitter, and Jesper Q. Svejstrup. 2020. “Using TTchem-Seq for Profiling Nascent Transcription and Measuring Transcript Elongation.” Nature Protocols 15 (2): 604–27.

Haberle, V., A. R. Forrest, Y. Hayashizaki, P. Carninci, and B. Lenhard. 2015. “CAGEr: Precise TSS Data Retrieval and High-Resolution Promoterome Mining for Integrative Analyses.” Nucleic Acids Research 43 (8): e51.

Harrison, Peter W., M. Ridwan Amode, Olanrewaju Austine-Orimoloye, Andrey G. Azov, Matthieu Barba, If Barnes, Arne Becker, et al. 2024. “Ensembl 2024.” Nucleic Acids Research 52 (D1): D891–99.

Henriques, Telmo, Benjamin S. Scruggs, Michiko O. Inouye, Ginger W. Muse, Lucy H. Williams, Adam B. Burkholder, Christopher A. Lavender, David C. Fargo, and Karen Adelman. 2018. “Widespread Transcriptional Pausing and Elongation Control at Enhancers.” Genes & Development 32 (1): 26–41.

Hirabayashi, Shigeki, Shruti Bhagat, Yu Matsuki, Yujiro Takegami, Takuya Uehata, Ai Kanemaru, Masayoshi Itoh, et al. 2019. “NET-CAGE Characterizes the Dynamics and Topology of Human Transcribed Cis-Regulatory Elements.” Nature Genetics 51 (9): 1369–79.

Kruesi, William S., Leighton J. Core, Colin T. Waters, John T. Lis, and Barbara J. Meyer. 2013. “Condensin Controls Recruitment of RNA Polymerase II to Achieve Nematode X-Chromosome Dosage Compensation.” ELife 2 (June): e00808.

Langmead, Ben, and Steven L. Salzberg. 2012. “Fast Gapped-Read Alignment with Bowtie 2.” Nature Methods 9 (4): 357–59.

Larke, Martin S. C., Ron Schwessinger, Takayuki Nojima, Jelena Telenius, Robert A. Beagrie, Damien J. Downes, A. Marieke Oudelaar, et al. 2021. “Enhancers Predominantly Regulate Gene Expression during Differentiation via Transcription Initiation.” Molecular Cell 81 (5): 983–997.e7.

Love, M. I., W. Huber, and S. Anders. 2014. “Moderated Estimation of Fold Change and Dispersion for RNA-Seq Data with DESeq2.” Genome Biology 15 (12): 550.

Lu, Zhaolian, and Zhenguo Lin. 2019. “Pervasive and Dynamic Transcription Initiation in Saccharomyces Cerevisiae.” Genome Research 29 (7): 1198–1210.

Luse, Donal S., Mrutyunjaya Parida, Benjamin M. Spector, Kyle A. Nilson, and David H. Price. 2020. “A Unified View of the Sequence and Functional Organization of the Human RNA Polymerase II Promoter.” Nucleic Acids Research 48 (14): 7767–85.

Mahat, Dig Bijay, Hojoong Kwak, Gregory T. Booth, Iris H. Jonkers, Charles G. Danko, Ravi K. Patel, Colin T. Waters, Katie Munson, Leighton J. Core, and John T. Lis. 2016. “Base-Pair-Resolution Genome-Wide Mapping of Active RNA Polymerases Using Precision Nuclear Run-on (PRO-Seq).” Nature Protocols 11 (8): 1455–76.

Malabat, Christophe, Frank Feuerbach, Laurence Ma, Cosmin Saveanu, and Alain Jacquier. 2015. “Quality Control of Transcription Start Site Selection by Nonsense-Mediated-MRNA Decay.” ELife 4 (April). 10.7554/eLife.06722.

Martin, Marcel. 2011. “Cutadapt Removes Adapter Sequences from High-Throughput Sequencing Reads.” EMBnet.Journal 17 (1): 10.

Nechaev, Sergei, David C. Fargo, Gilberto dos Santos, Liwen Liu, Yuan Gao, and Karen Adelman. 2010. “Global Analysis of Short RNAs Reveals Widespread Promoter-Proximal Stalling and Arrest of Pol II in Drosophila.” Science (New York, N.Y.) 327 (5963): 335–38.

Noguchi, Shuhei, Takahiro Arakawa, Shiro Fukuda, Masaaki Furuno, Akira Hasegawa, Fumi Hori, Sachi Ishikawa-Kato, et al. 2017. “FANTOM5 CAGE Profiles of Human and Mouse Samples.” Scientific Data 4 (1): 170112.

Pelechano, V., W. Wei, and L. M. Steinmetz. 2013. “Extensive Transcriptional Heterogeneity Revealed by Isoform Profiling.” Nature 497 (7447): 127–31.

Policastro, Robert A., Daniel J. McDonald, Volker P. Brendel, and Gabriel E. Zentner. 2021. “Flexible Analysis of TSS Mapping Data and Detection of TSS Shifts with TSRexploreR.” NAR Genomics and Bioinformatics 3 (2): lqab051.

Policastro, Robert A., R. Taylor Raborn, Volker P. Brendel, and Gabriel E. Zentner. 2020. “Simple and Efficient Profiling of Transcription Initiation and Transcript Levels with STRIPE-Seq.” Genome Research 30 (6): 910–23.

Quinlan, Aaron R., and Ira M. Hall. 2010. “BEDTools: A Flexible Suite of Utilities for Comparing Genomic Features.” Bioinformatics (Oxford, England) 26 (6): 841–42.

Sartorelli, Vittorio, and Shannon M. Lauberth. 2020. “Enhancer RNAs Are an Important Regulatory Layer of the Epigenome.” Nature Structural & Molecular Biology 27 (6): 521–28.

Schwalb, B., M. Michel, B. Zacher, K. Fruhauf, C. Demel, A. Tresch, J. Gagneur, and P. Cramer. 2016. “TT-Seq Maps the Human Transient Transcriptome.” Science 352 (6290): 1225–28.

Smith, Tom, Andreas Heger, and Ian Sudbery. 2017. “UMI-Tools: Modeling Sequencing Errors in Unique Molecular Identifiers to Improve Quantification Accuracy.” Genome Research 27 (3): 491–99.

Sousa-Luis, R., G. Dujardin, I. Zukher, H. Kimura, C. Weldon, M. Carmo-Fonseca, N. J. Proudfoot, and T. Nojima. 2021. “POINT Technology Illuminates the Processing of Polymerase-Associated Intact Nascent Transcripts.” Molecular Cell. 10.1016/j.molcel.2021.02.034.

Takahashi, Hazuki, Sachi Kato, Mitsuyoshi Murata, and Piero Carninci. 2012. “CAGE (Cap Analysis of Gene Expression): A Protocol for the Detection of Promoter and Transcriptional Networks.” Methods in Molecular Biology (Clifton, N.J.) 786: 181–200.

Thorsen, Kasper, Troels Schepeler, Bodil Øster, Mads H. Rasmussen, Søren Vang, Kai Wang, Kristian Q. Hansen, et al. 2011. “Tumor-Specific Usage of Alternative Transcription Start Sites in Colorectal Cancer Identified by Genome-Wide Exon Array Analysis.” BMC Genomics 12 (October): 505.

Vo Ngoc, Long, California Jack Cassidy, Cassidy Yunjing Huang, Sascha H. C. Duttke, and James T. Kadonaga. 2017. “The Human Initiator Is a Distinct and Abundant Element That Is Precisely Positioned in Focused Core Promoters.” Genes & Development 31 (1): 6–11.

Wang, E. T., R. Sandberg, S. Luo, I. Khrebtukova, L. Zhang, C. Mayr, S. F. Kingsmore, G. P. Schroth, and C. B. Burge. 2008. “Alternative Isoform Regulation in Human Tissue Transcriptomes.” Nature 456 (7221): 470–76.

Wilkins, Oscar G., Charlotte Capitanchik, Nicholas M. Luscombe, and Jernej Ule. 2021. “Ultraplex: A Rapid, Flexible, All-in-One Fastq Demultiplexer.” Wellcome Open Research 6 (June): 141.

