## Supplementary material for "Mapping and quantifying nascent transcript start sites using TT-TSS-seq": Figures S1 S2

Figure S1

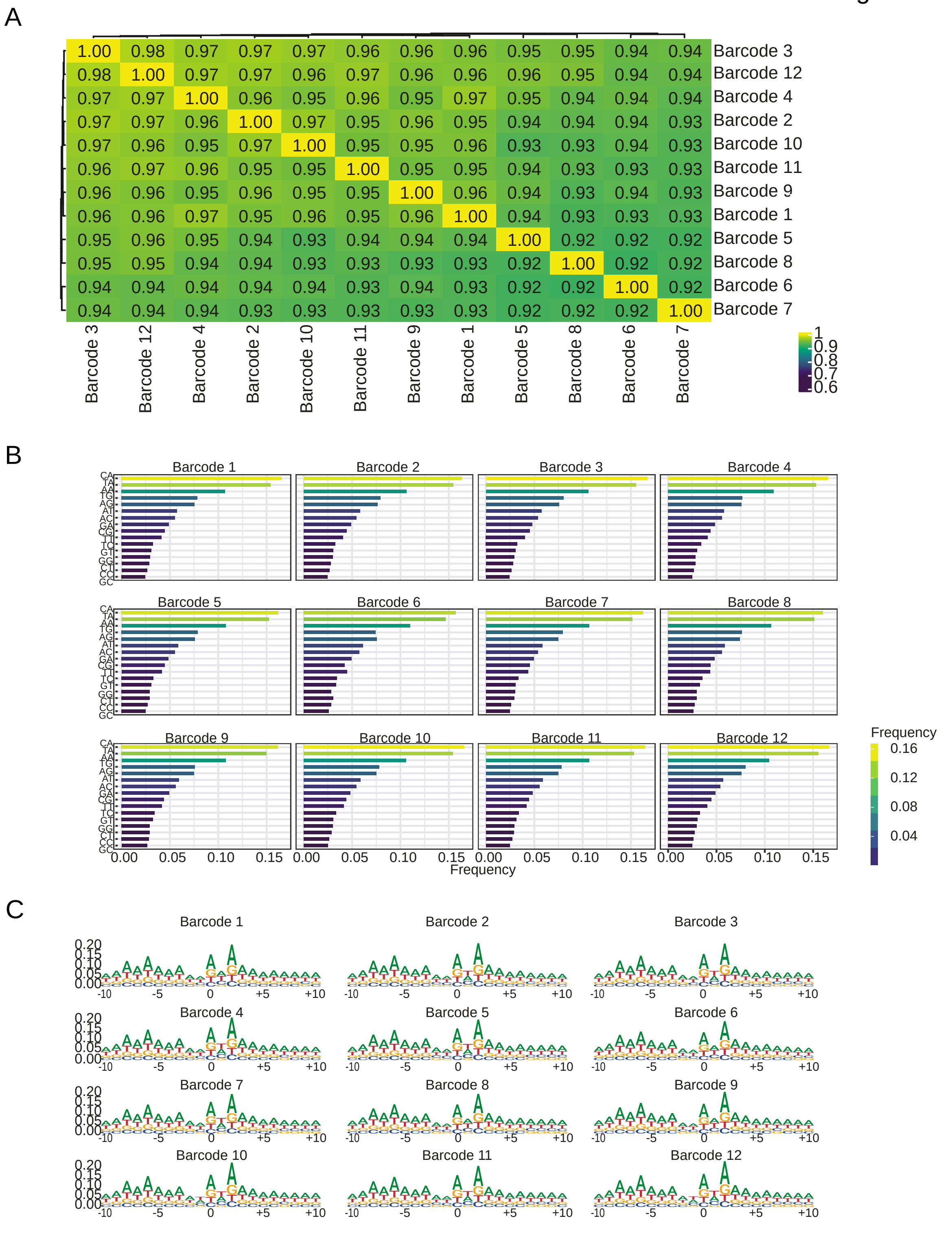


**Figure S1.** Testing of sample multiplexing in optimized TSS-seq protocol.
**(A)** Hierarchically clustered heatmap of Pearson's correlation coefficients (r values) between the determined TSSs of the libraries produced using the different 5' adaptor strategies. **(B)** Dinucleotide frequency analysis. The first nucleotide represents the -1 nucleotide and the second nucleotide represents the +1 tag/TSS identified by polyA-TSS-seq. **(C)** Sequence logos showing ±10 nucleotides surrounding the detected TSS (position 0). Analysis performed using TSRexploreR.

Figure S2

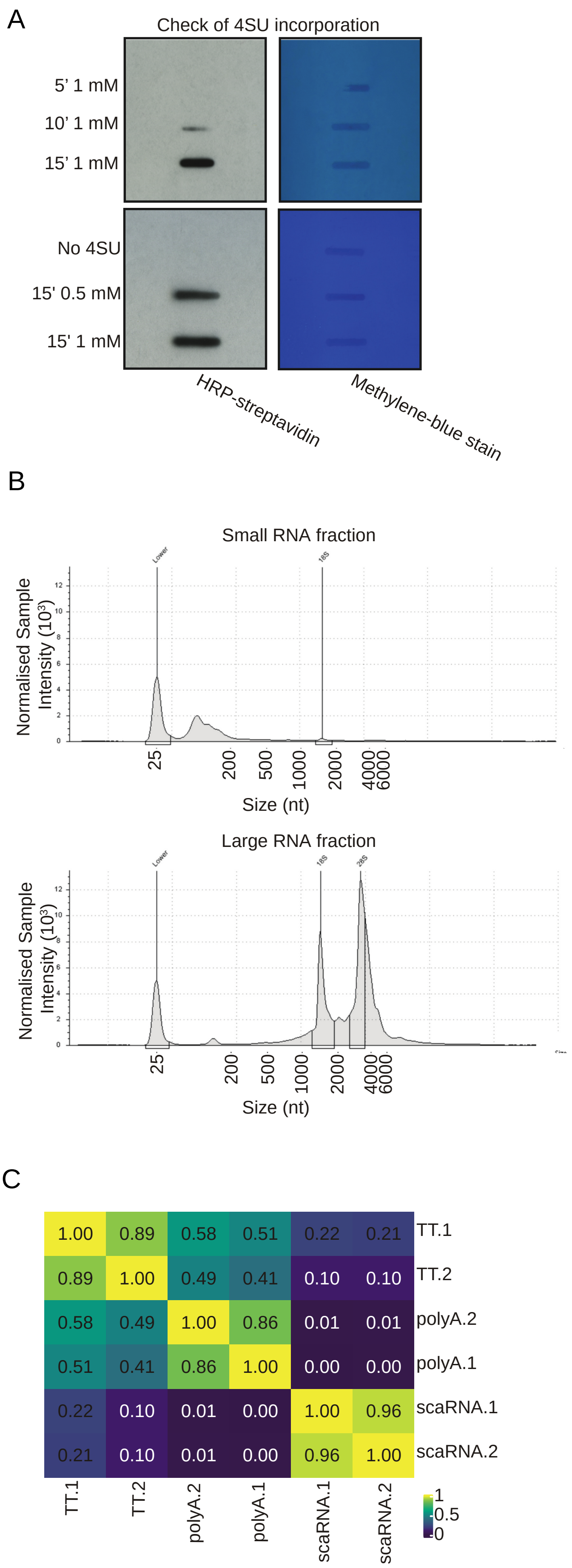
A
Check of 4SU incorporation
5’ 1 mM
10’ 1 mM
15’ 1 mM
No 4SU
15' 0.5 mM
15' 1 mM
HRP-streptavidin
Methylene-blue stain
B
Small RNA fraction
Normalised Sample Intensity (10³)
25
200
500
1000
2000
4000
6000
Size (nt)
Large RNA fraction
Normalised Sample Intensity (10³)
25
200
500
1000
2000
4000
6000
Size (nt)
C
1.00 0.89 0.58 0.51 0.22 0.21 TT.1
0.89 1.00 0.49 0.41 0.10 0.10 TT.2
0.58 0.49 1.00 0.86 0.01 0.01 polyA.2
0.51 0.41 0.86 1.00 0.00 0.00 polyA.1
0.22 0.10 0.01 0.00 1.00 0.96 scaRNA.1
0.21 0.10 0.01 0.00 0.96 1.00 scaRNA.2
TT.1
TT.2
polyA.2
polyA.1
scaRNA.1
scaRNA.2
1
0.5
0

**Figure S2.** TT-TSS-seq protocol development and comparisons.
**(A)** Assessment of 4SU incorporation in mESCs using slot blot, using different labelling times and concentrations. 4SU-containing RNA was labelled using a biotin linker, and HRP-conjugated streptavidin. Methylene blue staining was used to assess RNA loading levels. **(B)** RNA electropherogram, as measured using TapeStation, showing the sizes of RNA in the flow-through and eluate after size selection. A 3:1 ratio of RNA to ethanol was used. The 25 nt peak represents the ladder. **(C)** Hierarchically clustered heatmap of Pearson's correlation coefficients (r values) between the determined TSRs of the different samples. Tag counts were normalised using the DESeq2 median-of-ratios approach using a threshold of three counts. Tags within 25 bp, and to a maximum distance of 250 bp, were merged to identify TSRs.
