## Supplementary material for "Mapping and quantifying nascent transcript start sites using TT-TSS-seq": Table S1

| **Reaction** | **Original conditions** | **Optimised conditions** |
| --- | --- | --- |
| Phosphatase reaction | 2.4 U rSAP/ µg RNA  2 hours, 37°C | 2.4 U quickCIP/ µg RNA  2 hours, 37°C |
| Decapping reaction | 0.6 U CAPCLIP/ µg RNA  2 hours, 37°C | 1.2 U MDE/ µg RNA  2 hours, 37°C |
| 5’ adaptor ligation | 0% PEG  Overnight, 16°C | 5% PEG  RNA + adaptor heat denatured  2 hours 25°C, overnight 16°C |
| 3’ adaptor ligation | 10% PEG  2 hours 25°C, overnight 16°C | 15% PEG  RNA + adaptor heat denatured  2 hours 25°C, overnight 16°C |

Table S1. Summary TSS-seq enzymatic reaction conditions.

For the phosphatase reaction, rSAP was changed to quickCIP. For the decapping reaction, CAPCLIP was changed to MDE. A heat denaturation step was introduced before both adaptor ligation steps, and the concentration of PEG was also optimised. The 5’ adaptor reaction conditions were also altered.
